# Associations between impulsivity, risk behavior and HIV, HBV, HCV and syphilis seroprevalence among female prisoners in Indonesia: A cross-sectional study

**DOI:** 10.1101/468694

**Authors:** Rachel M. Arends, Erni J. Nelwan, Ratna Soediro, Reinout van Crevel, Bachti Alisjahbana, Herdiman T. Pohan, A. Katinka L. von Borries, Aart H. Schene, André J. A. M. van der Ven, Arnt F. A. Schellekens

## Abstract

HIV, hepatitis B and C, and syphilis share common transmission routes of which primarily unsafe sexual contact and injecting drug use are important. Impulsivity is a major factor contributing to this transmission risk behavior, however comprehensive studies within female, prison, and Asian populations are scarce. This cross-sectional study aims to delineate the contributions of different aspects of impulsivity to transmission risk behavior, among female inmates living in a prison in Jakarta (N=214). The relationships between various aspects of impulsivity, risky behavior and seropositivity were tested using analyses of variance and logistic regression analyses. Motor impulsivity was related to alcohol use, reward-related impulsivity to drug use, and cognitive/goal-directed impulsivity to sexual risk behavior. Finally, goal-directed impulsivity was also directly associated with seropositivity. Specific aspects of impulsivity are associated with different types of risky behaviors in Indonesian female prisoners, what can be relevant for future studies on infection prevention strategies for such a population.

## Introduction

HIV, hepatitis B (HBV) and C (HCV), and syphilis pose a great global health burden [1, 2]. Global Burden of Disease (GBD) studies have shown that the total number of people living with HIV has been steadily increasing and reached around 36,9 million in 2017 [3]. This can be explained by rising epidemics in some parts of the world, like in Asia and the Pacific [3]. Indonesia is among the most afflicted countries in this region, with some of the highest numbers of new infections and people living with HIV [4].

The prevalence of syphilis and hepatitis also remain high in key populations across the globe. With globally around 6 million new cases of syphilis infections, syphilis still poses a major health burden [2]. Moreover, viral hepatitis remains a leading cause of death and disability-adjusted life years (DALYs) worldwide, with the greatest burden in south-east Asia [5]. As such, these infections as HIV, HBV, HCV and syphilis have a serious impact on sexual and public health across the globe, particularly in Southeast Asian countries like Indonesia [6].

HIV, HBV, HCV and syphilis share common ways of transmission through blood or other body fluids, mainly as a result of unsafe sexual intercourse or intravenous drug use (IDU) [6]. In Southeast Asia 21% of the burden of HIV cases and 28% of HCV were attributable to IDU in 2013, frequently found among populations such as sex workers, men who have sex with men (MSM) or inmates [1].

General prevention programmes often fail to reach these specific key populations and the decline in those HIV infections has slowed down in recent years, also among inmates [7]. The prevalence of HIV infection in 2016 was estimated to be five times higher in prison populations and 24 times higher when inmates inject drugs, compared to the general population [7], possibly due to persisting or increasing risks such as drug use, rape and unsafe sex [8]. As 30 million people spend time in prisons every year, it is crucial to unravel factors associated with persistent transmission risk behavior in inmates, in order to target the ongoing epidemics of HIV, HBV, HCV, and syphilis [8].

A major determinant of risky behavior is impulsivity. People who are more impulsive can e.g. show more negative life outcomes such as criminal activities, problems with substance use, and incidence of STIs [9, 10]. Impulsivity is a broad construct indicating a tendency to act without deliberate thinking, reflection, or consideration of the consequences [9]. It includes a number of independent components, such as impaired inhibitory motor or cognitive control, preference for immediate rewards and goal-directed or sensation seeking behavior [11].

Because impulsivity is associated with various risk-taking behaviors and therefore increased probabilities of contracting infections [9-11], impulsivity represents an important construct contributing to public health concerns. Previous studies showed that specific aspects of impulsivity may contribute to different types of risky behaviors. For example, goal-directed impulsivity has been associated with sexual risk behavior among HIV-positive persons [12], while e.g. motor impulsivity has been associated with substance use and sharing drug equipment [13]. A major limitation of these studies is that they take only one aspect of impulsivity in consideration in the context of a single type of risky behavior.

Another major issue with current studies into the relationship between risky behavior and impulsivity in infection key populations, is that they are mostly conducted among male, Western populations. Yet, in recent years female prisoners have contributed as much as or more than males to the transmission of HIV and co-infections [8]. Overall numbers of female patients are increasing and they seem to acquire HIV at a younger age compared to males [14]. Finally, associations between risky behaviors like substance abuse and unsafe sexual practices seem to be increased among women [15].

In this study we investigate the role of different aspects of impulsivity in various types of transmission risk behavior in female prisoners in Indonesia. Specifically, we achieved to examine if: a) goal-directed, cognitive and motor impulsivity are associated with sexual and/or substance use related risk behavior, and if b) the associations between different types of impulsivity and seropositivity for HIV, HBV, HCV and syphilis are mediated by distinct types of risk behavior.

## Materials and methods

### Design

To investigate the specific role of impulsivity in transmission risk behavior, this cross-sectional bio-behavioral study was conducted, combining self-reported behavioral data with a serosurvey on the presence of infectious diseases in the Jakarta Pondok Bambu women’s penitentiary.

### Setting & participants

All new incoming prisoners at the time of the study (N = 300) were offered serological testing (for diagnosis of HIV, HBV, HCV and/or syphilis). Prisoners participating in the serosurvey were also asked to participate in the behavioral part of the study, consisting of 6 self-report questionnaires. Two Indonesian field physicians and two Dutch psychology students were available during the data collection to explain the procedure and answer questions. Finally, of the 300 women 56 were untraceable or declined participation and 30 were excluded due to a lack of comprehension of the questionnaires e.g. because of illiteracy. In total 214 participants completed the research incl. serology testing and answering the six questionnaires (Fig 1).

**Fig 1.**
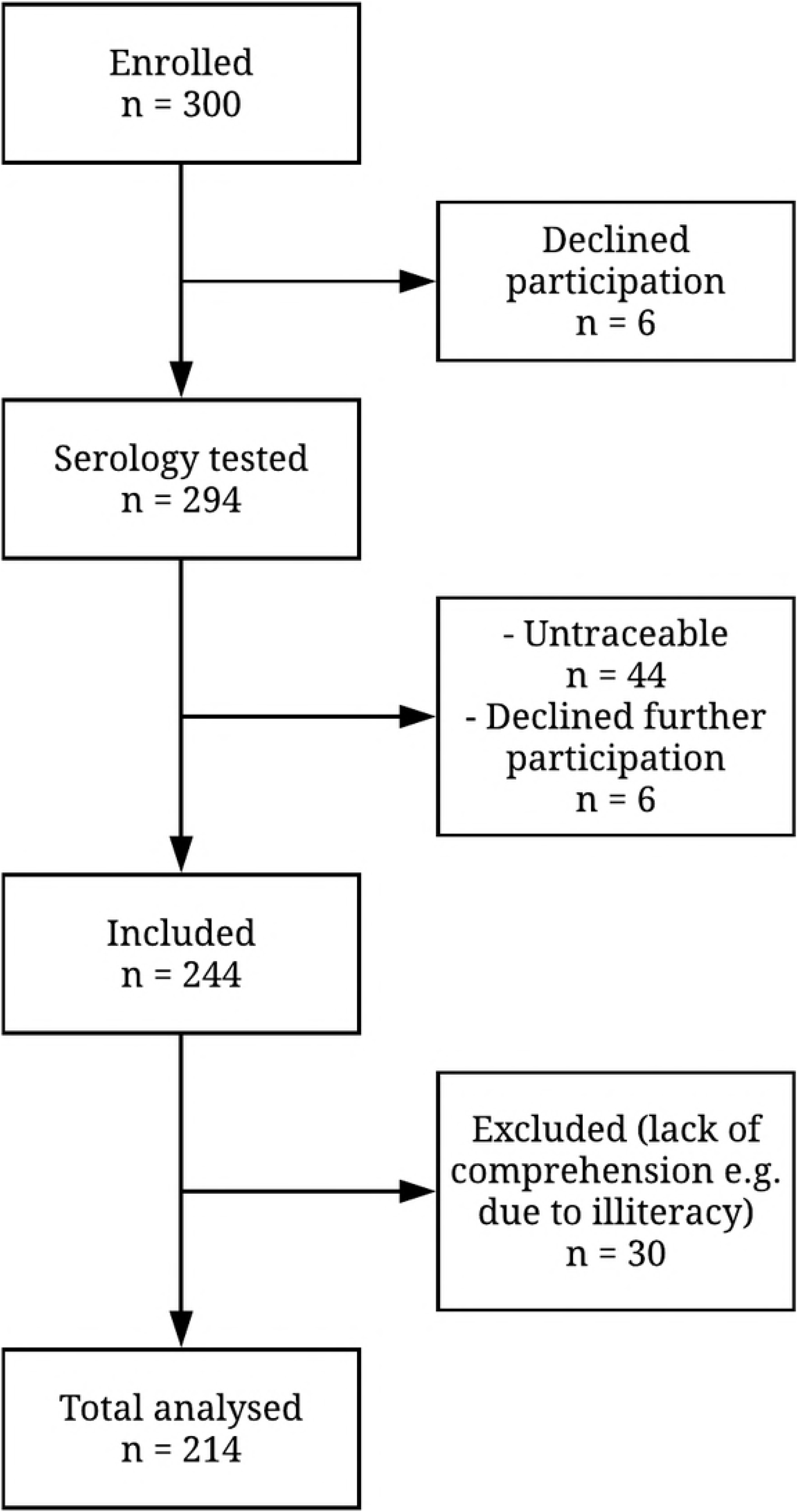
Study inclusion flow chart.

## Measurements

### Demographics

Demographic variables including age, education, marital status and occupation, were assessed using a short self-report questionnaire.

### Assessment of impulsivity

*The Barratt Impulsiveness Scale (BIS-11)* was used to assess cognitive and motor impulsivity [16]. It consists of 30 items, scoring on a four-point Likert scale, covering 3 domains of impulsivity. Motor impulsivity: BIS-motor (acting without thinking, 11 items), and two cognitive subscales: BIS-attentional (quick decision-making, 8 items), and BIS-non-planning (lack of forethought, 11 items). The English version of the BIS-11 has good internal consistency and retest reliability [16].

*The Behavioral Inhibition System and Behavioral Activation System (BIS/BAS)* was used to assess two self-regulatory systems, one of which bears on appetitive motivation, the other on aversive motivation [17]. It consists of 20 items, scored on a four-point Likert scale measuring self-control: the ability to regulate emotions, thoughts, and behavior in the face of temptations and impulses in order to achieve specific goals. The BIS-scale measures neuroticism and negative effect (avoidance tendencies), consisting of 7 items about regulating aversive motives. The BAS-scale measures extraversion and positive effect (approach tendencies) and consists of 3 subscales: BAS-drive (goal-directed impulsivity, 4 items), BAS-fun-seeking (sensation seeking-impulsivity, 4 items) and BAS-reward (reward-impulsivity, 5 items). The English version of the BIS/BAS has adequate psychometric properties [17].

*The Sensitivity to Punishment and Sensitivity to Reward Questionnaire (SPSRQ)* was used to assess reward-related impulsivity [18]. It consists of 48 items (yes/no) and measures punishment and reward responsiveness. Half of the items are summed in an index of ‘sensitivity to punishment’ and half as an index of ‘sensitivity to reward’. The English version of the SPSRQ is a reliable measure, independent of personality factors [18].

The questionnaires concerning impulsivity were not available in Indonesian language. These were first translated from the English versions into Indonesian, by two native Indonesian speakers fluent in English. This was followed by a back-translation into English by an English speaker, also fluent in Indonesian language (see acknowledgements). Any discrepancies were discussed and solved in a consensus meeting with all translators and executive researchers, in line with the WHO guidelines [19].

### Assessment of risky behaviors

*The Alcohol Use Disorders Identification Test* (AUDIT) is a method developed by the World Health Organization (WHO) to screen for excessive drinking patterns [20]. It consists of 10 items with a five-point Likert scale. The first 3 items cover the consumption pattern, the other 7 assess alcohol dependence and the consequences of alcohol use. For women a score ≥ 8 indicates harmful alcohol use. The AUDIT has excellent psychometric properties and has been used in different populations worldwide [20].

*The Drug Abuse Screening Test* (DAST) is a self-report questionnaire including 10 dichotomous items (yes/no), used to screen for problematic drug use (scores ≥ 3) [21]. The reliability of this commonly used instrument is shown in various populations [21].

*The Sexual Risk Behavior* (SRB) questionnaire was used to assess sexual risk behavior [22]. It entails 20 items regarding (risky) sexual behavior e.g. condom use and amount of sexual partners [22]. Questions not regarding risky behavior or applying only to male respondents were excluded, leaving 16 questions. Three items have a four-point scale, 5 items have a six-point scale, and 8 items have binary outcomes (yes/no).

The tree questionnaires concerning risky behavior were available in Indonesian language. Although reliability data of these translated versions were not available, we used these measurements based on mainly good psychometric characteristics of the English versions.

### Serosurvey

Screening for HIV, HBV, HCV and syphilis was done by serology testing. Screening for HIV was done by chemiluminescent magnetic microparticle immunoassay (CMIA), a rapid diagnostic HIV-1/2 test (PT Oncoprobe Utama), and confirmed by enzyme-linked fluorescence assay (ELFA, VIDAS^®^ HIV DUO ULTRA). HBV serology was performed by using CMIA method for hepatitis B surface antigen (HBsAg) (using ARCHITECT HBsAg qualitative assay and ARCHITECT HBsAg Qualitative Confirmatory Assay by ABBOTT Diagnostics^®^) and antibody hepatitis B core antigen (Anti-HBc) (using ARCHITECT Anti-HBc IgM assay and ARCHITECT Anti-HBc II assay by ABBOTT Diagnostics^®^). Active HBV infection was diagnosed if HBsAg and anti-HBc were positive. HCV was determined by using in vitro diagnostic immunoassay for the detection of immunoglobulin G (IgG) antibodies to Hepatitis C Virus (using ADVIA Centaur HCV assay). Finally, syphilis was tested with Rapid Plasma Reagin (RPR) test (using IMMUTREP RPR by Omega Diagnostics^®^); positive result of RPR test for syphilis was then confirmed by TPHA (Treponema pallidum haemagglutination: solid phase immunochromatographic assay for the qualitative detection of antibodies of all isotypes (IgG, IgM, IgA) against *Treponema pallidum* (TP) by SD Bioline Syphillis 3.0 Multi by Standard Diagnostics, Inc^®^).

### Data-analysis

First, demographic variables (age, education, occupation, marital status, committed crime) and infection rates (one or multiple infections) were analyzed. An univariate Analysis of Variance (ANOVA) was used to compare the continuous variable age, the other four categorical variables were examined using Chi-square tests.

Next, subscale scores and total scores were calculated for all questionnaires. For the three impulsivity measurements this included nine subscales (BIS-11, BIS/BAS and SPSRQ subscales). Two subscales were not included in the analyses, as these did not measure impulsivity (BIS scale of the BIS/BAS and the punishment scale of the SPSRQ), leaving seven continuous impulsivity variables. Three total scores were calculated for the three types of risky behavior, which were transformed into categorical variables (low vs. high risky behavior) according to established cut-off scores (alcohol-: AUDIT > 7, drugs-: DAST > 2, and sex-related: SRB > 0).

Finally, to test the hypotheses, binary and multinomial logistic regression analyses were conducted, using 5000 bootstrap samples and enter method. The first hypothesis, that different aspects of impulsivity are associated with various types of risky behaviors, was tested using three separate logistic regression analyses for each form of (alcohol-, drugs, and sex-related) risky behavior. The impulsivity variables were tested for multicollinearity. Moreover, age was included in the model, because risk behavior has shown to be associated with age [23]. The second hypothesis, that a relationship between impulsivity and the prevalence of viral infections can be mediated by risky behavior, was tested using the significant impulsivity factors of the first analyses in a final logistic regression analysis.

## Results

### Descriptive statistics

Participants with risky behavior (alcohol-, drugs- or sex-related) were younger (*F*(1, 214) = 5.77, *p* = .017) and were more often convicted for drug-related crimes, compared to those without risky behavior (χ^2^(1) = 23.56, *p* < .001). Other demographic variables did not differ between participants with and without risky behavior (*p* > .069). Infectious diseases were diagnosed in 38 inmates (17.8%), including HIV (3.7%), syphilis (7.0%), HBV (3.3%), and HCV (2.3%). Detailed study characteristics are shown in Table 1.

**Table 1.** Demographic and medical characteristics of female inmates with and without transmission risk behavior.

|  | Total group |  | Risk behavior |  | No risk behavior |  | Pearson | *p* |
| --- | --- | --- | --- | --- | --- | --- | --- | --- |
| | *n*<br>(214) | % | *n*<br>(144) | % | *n*<br>(70) | % | $\chi^2$ | |
| Demographics |  |  |  |  |  |  |  |  |
| Education<sup>a</sup> |  |  |  |  |  |  | 3.217 | .200 |
| no education | 11 | 5.1 | 5 | 3.5 | 6 | 8.6 |  |  |
| low education | 77 | 36.0 | 50 | 34.7 | 27 | 38.6 |  |  |
| high education | 126 | 58.9 | 89 | 61.8 | 37 | 52.9 |  |  |
| Occupation |  |  |  |  |  |  | 1.297 | .523 |
| unemployed | 47 | 22.0 | 29 | 20.1 | 18 | 25.7 |  |  |
| employee/entrepreneur | 166 | 77.6 | 114 | 79.2 | 52 | 74.3 |  |  |
| Marital status |  |  |  |  |  |  | 5.356 | .069 |
| married | 120 | 56.1 | 73 | 50.7 | 47 | 67.1 |  |  |
| divorced | 61 | 28.5 | 24 | 16.7 | 14 | 20.0 |  |  |
| single | 33 | 15.4 | 47 | 32.6 | 9 | 12.9 |  |  |
| Crime |  |  |  |  |  |  | 23.562 | <.001* |
| drug-related | 109 | 50.9 | 90 | 62.5 | 19 | 27.1 |  |  |
| other | 105 | 49.1 | 54 | 37.5 | 51 | 72.9 |  |  |
| Serology |  |  |  |  |  |  | 8.344 | .138 |
| HIV | 8 | 3.7 | 7 | 4.9 | 1 | 1.4 |  |  |
| HBV | 15 | 3.3 | 8 | 5.6 | 7 | 10.0 |  |  |
| HCV | 7 | 2.3 | 7 | 4.9 | 0 | 0.0 |  |  |
| syphilis | 5 | 7.0 | 4 | 2.8 | 1 | 1.4 |  |  |
| multiple infections<sup>b</sup> | 3 | 1.4 | 3 | 2.1 | 0 | 0.0 |  |  |
|  | *M* | *SD* | *M* | *SD* | *M* | *SD* | *F* | *p* |
| Age (years) | 33.39 | 9.87 | 32.27 | 9.29 | 36.81 | 10.19 | 5.768 | .017* |

\**p* < .05
<sup>a</sup> Low education: elementary school and junior high school, high education: high school and university
<sup>b</sup> Having a combination of infections, in this sample found: HBV-syphilis; HCV-HIV; HBV-HCV-HIV

### Associations between impulsivity and risky behavior

The first set of regression analyses showed a significant prediction model for problematic alcohol use, based on impulsivity measures (χ^2^(8) = 30.95, *p* < .001, Nagelkerke R^2^ = .20), with an overall correct prediction of 76%. The Wald criterion demonstrated that BIS-Motor impulsivity (*p* = .016) and age (*p* = .003) significantly contributed to the prediction. Similarly, impulsivity reliably predicted problematic drug use (χ^2^(8) = 23.99, *p* = .002, Nagelkerke R^2^ = .15). The overall correct prediction of the model was 71%, with SPSRQ-reward significantly contributing to the prediction (*p* = .016). Impulsivity measures also predicted sexual risk behavior (χ^2^(8) = 22.64, *p* = .004, Nagelkerke R^2^ = .13). The overall correct prediction of the model was 65%. BIS-attentional (*p* = .012) and BAS-drive (*p* = .034) made a significant contribution to the prediction (Table 2).

**Table 2.**
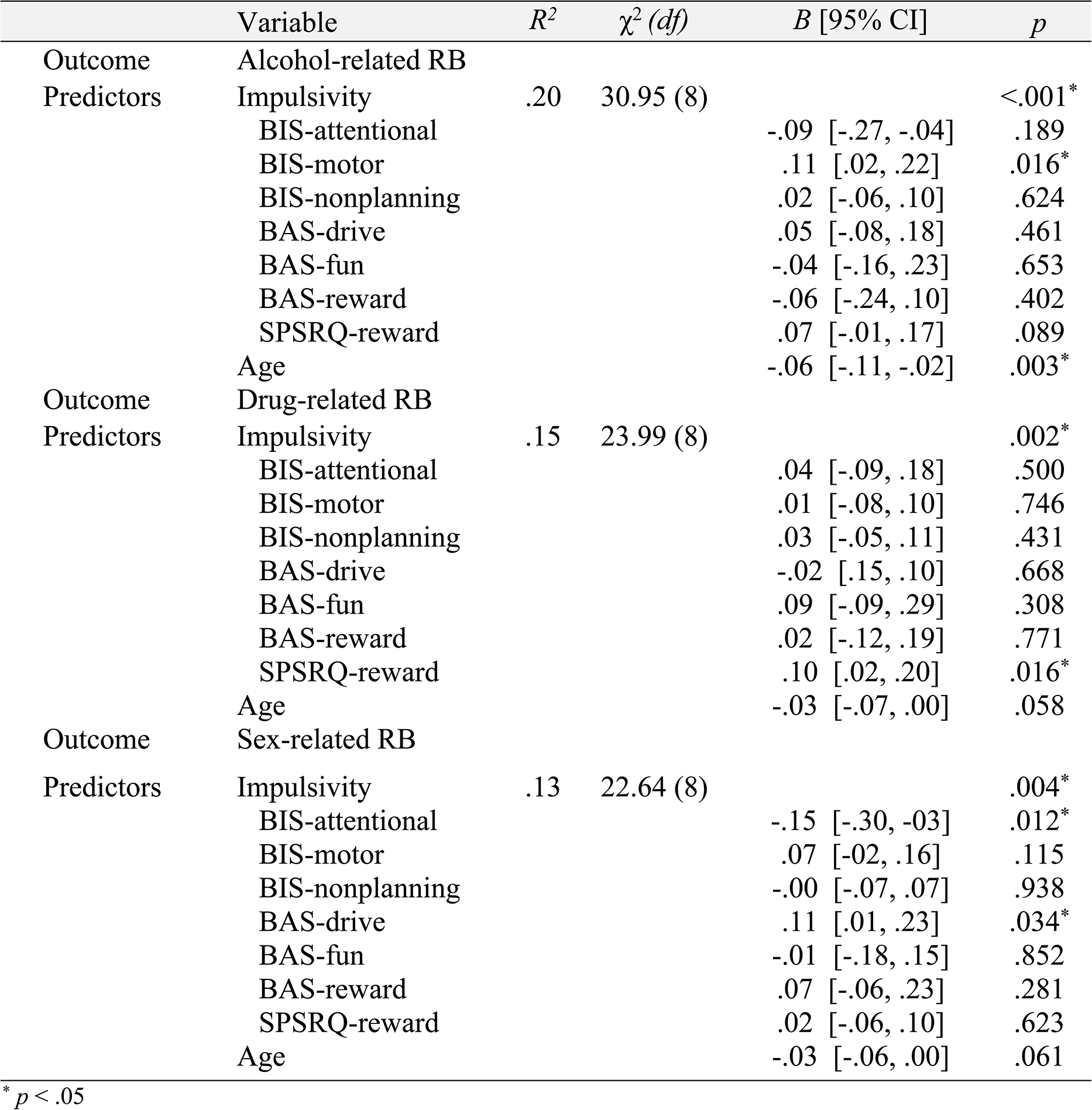
Associations between impulsivity and transmission risk behavior among female inmates.

| | Variable | $R^2$ | $\chi^2 (df)$ | $B$ [95% CI] | $p$ |
| --- | --- | --- | --- | --- | --- |
| Outcome | Alcohol-related RB |  |  |  |  |
| Predictors | Impulsivity | .20 | 30.95 (8) |  | <.001* |
|  | BIS-attentional |  |  | -.09 [-.27, -.04] | .189 |
|  | BIS-motor |  |  | .11 [.02, .22] | .016* |
|  | BIS-nonplanning |  |  | .02 [-.06, .10] | .624 |
|  | BAS-drive |  |  | .05 [-.08, .18] | .461 |
|  | BAS-fun |  |  | -.04 [-.16, .23] | .653 |
|  | BAS-reward |  |  | -.06 [-.24, .10] | .402 |
|  | SPSRQ-reward |  |  | .07 [-.01, .17] | .089 |
|  | Age |  |  | -.06 [-.11, -.02] | .003* |
| Outcome | Drug-related RB |  |  |  |  |
| Predictors | Impulsivity | .15 | 23.99 (8) |  | .002* |
|  | BIS-attentional |  |  | .04 [-.09, .18] | .500 |
|  | BIS-motor |  |  | .01 [-.08, .10] | .746 |
|  | BIS-nonplanning |  |  | .03 [-.05, .11] | .431 |
|  | BAS-drive |  |  | -.02 [-.15, .10] | .668 |
|  | BAS-fun |  |  | .09 [-.09, .29] | .308 |
|  | BAS-reward |  |  | .02 [-.12, .19] | .771 |
|  | SPSRQ-reward |  |  | .10 [.02, .20] | .016* |
|  | Age |  |  | -.03 [-.07, .00] | .058 |
| Outcome | Sex-related RB |  |  |  |  |
| Predictors | Impulsivity | .13 | 22.64 (8) |  | .004* |
|  | BIS-attentional |  |  | -.15 [-.30, -.03] | .012* |
|  | BIS-motor |  |  | .07 [-.02, .16] | .115 |
|  | BIS-nonplanning |  |  | -.00 [-.07, .07] | .938 |
|  | BAS-drive |  |  | .11 [.01, .23] | .034* |
|  | BAS-fun |  |  | -.01 [-.18, .15] | .852 |
|  | BAS-reward |  |  | .07 [-.06, .23] | .281 |
|  | SPSRQ-reward |  |  | .02 [-.06, .10] | .623 |
|  | Age |  |  | -.03 [-.06, .00] | .061 |

\* $p < .05$

### Mediation of impulsivity on infections by risk behavior

The final regression analysis showed a significant prediction model for infectious diseases based on impulsivity measures (χ^2^(5) = 14.39, *p* = .013, Nagelkerke R^2^ = .11). The overall correct prediction of the model was 82%. According to the Wald criterion, BAS-drive impulsivity significantly contributed to the prediction (*p* = .017). When adding risk behavior as predictors, the model became less significant (χ^2^(8) = 18.29, *p* = .019, Nagelkerke partial R^2^ = .14) but the total overall correct prediction did not change (82%). So, the relationship of impulsivity and the viral infections could not be explained by a mediation effect of risk behavior (χ^2^(3) = 4.20, *p* = .241, Nagelkerke R^2^ = .03). See Table 3 and Fig 2 for further details.

**Table 3.**
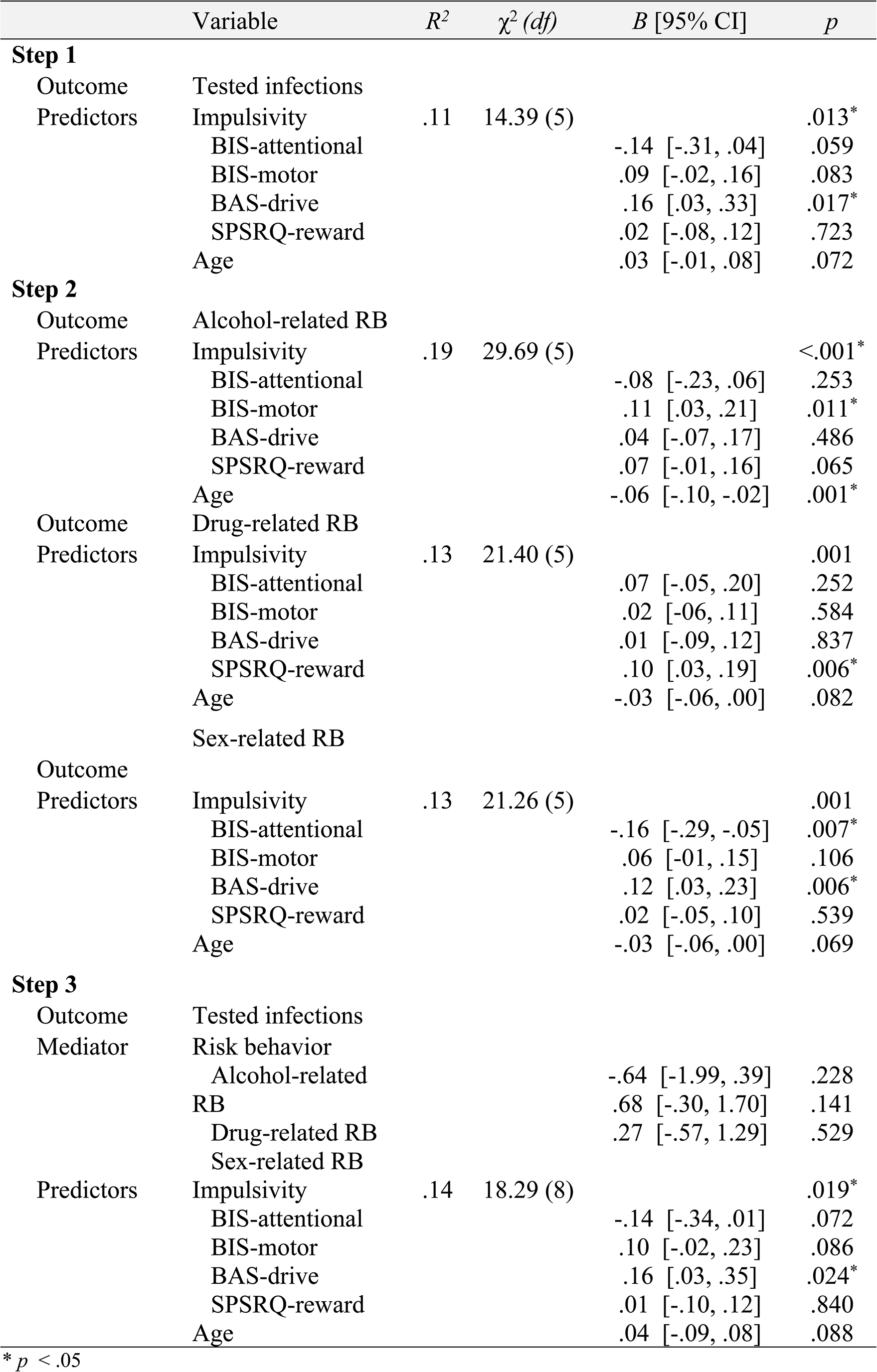
Associations between impulsivity, transmission risk behavior (hypothesis 1, step 2) and infections HIV, hepatitis B and C and syphilis (hypothesis 2, step 1 and 3) among female inmates.

**Fig 2.**
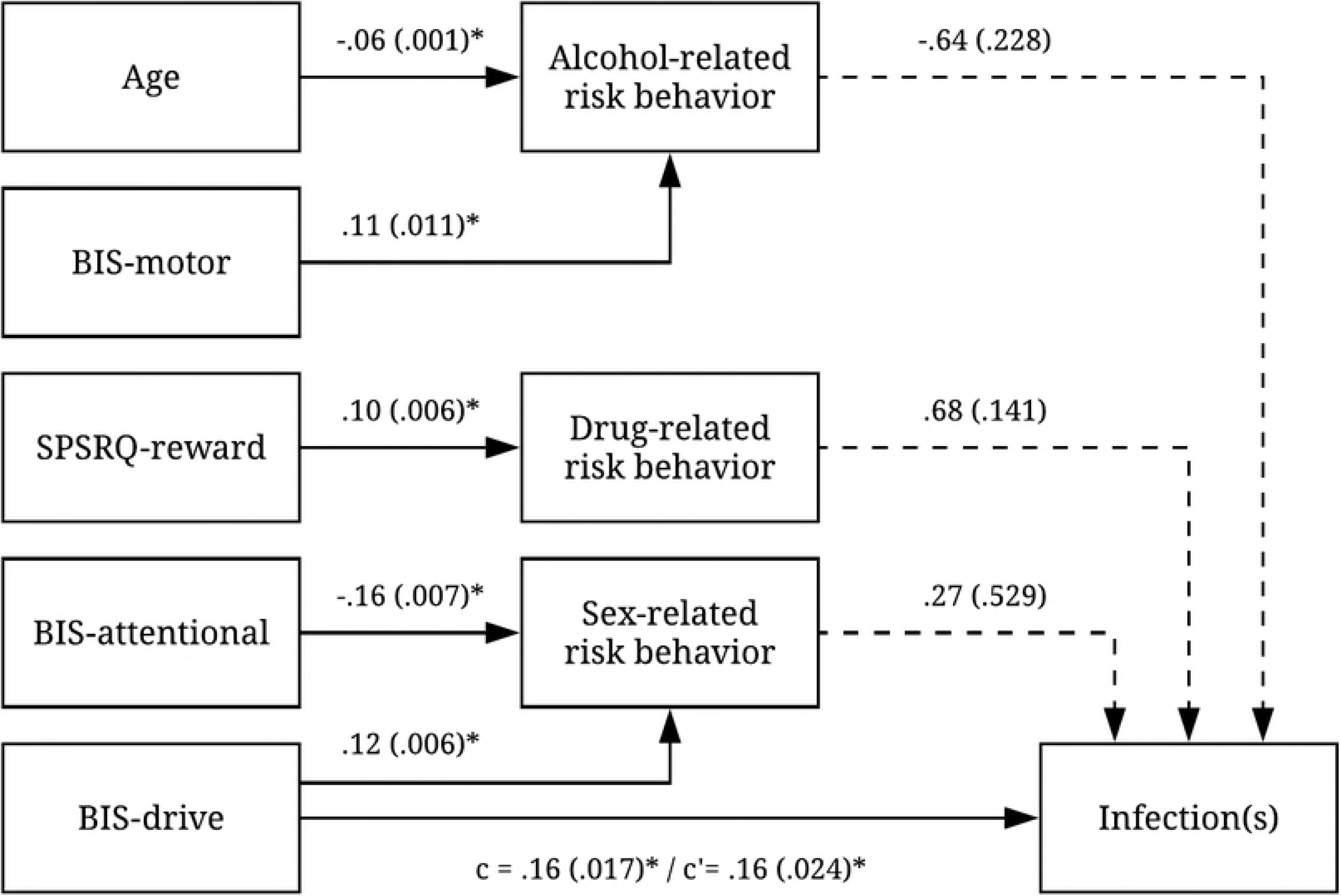
Associations of impulsivity and age with risk behaviors and viral infections. * *p* < .05

## Discussion

The results found in this study are in line with research showing a relationship between impulsivity and substance use or the incidence of viral infections such as HIV, HBV, HCV and syphilis [9, 10]. Furthermore, do these findings confirm prior research on the relationship between various aspects of impulsivity and risky behaviors [9, 24]. Interestingly, sexual risk behavior has become an increasingly important risk factor for the above mentioned infections in South-East Asia [25]. Our finding that goal-directed impulsivity was related with sexual risk behavior, and seropositivity for HIV, HBV, HCV or syphilis might clarify these observations.

One of the few studies that studied the subject of impulsivity and risk behavior in the context of HIV infection, distinguished two types of impulsivity: hot and cool [26]. Hot impulsivity concerns affectively-mediated preferences for immediate gratification in the presence of anticipatory cues (e.g. reward and goal-related impulsivity). Cool impulsivity involves a more affectively neutral tendency towards rapid, premature responses (such as motoric and attentional impulsivity). In line with our finding that goal-directed behavior is related to sexual risk behavior and seropositivity for HIV, HBV, HCV or syphilis, they found that especially hot dimensions were related to sexual risk behaviors [26]. Our results also confirm the previously suggested association between motor impulsivity (or cool impulsivity) and alcohol use [27]. These observations also affirm the hypothesis that there might be separate and independent processes involved in the relationship between impulsivity and risky behavior.

It is unclear whether distinctive brain areas are involved in these different aspects of impulsivity and risky behavior. Different forms of impulsivity have been associated with specific brain circuits [26]. For instance, reward impulsivity has been associated with alterations within the ventromedial orbitofrontal-limbic brain circuits regulating motivation and cognitive impulsivity has been associated with alterations within more dorsolateral prefrontal cortex areas [28]. Furthermore, based on our findings it is impossible to delineate state and trait related factors contributing the different types of impulsivity. Which genetic and developmental trait influences and which state dependent influences of infection or medication side effects might be involved, can be a focus of future research.

The relevance of the current findings for prevention strategies for transmission of HIV, hepatitis B and C and syphilis remains to be explored. As risky behaviors often continue or even increase in prison settings [29], the current findings warrant further studies into the potential of impulse-control or self-regulation programs to target risky behavior. For instance mindfulness training and behavioral training strategies have been shown to successfully alter impulsivity and risky behavior in several high impulsive populations [30].

The current findings should be seen in the light of some study limitations. Psychometric information on the Indonesian versions of the questionnaires that were used is lacking, and only self-report measurements for impulsivity factors and risky behaviors were used. This might have reduced validity of the results that we found. Future studies should perform psychometric evaluation of the Indonesian versions of the applied questionnaires and add behavioral tasks to have more direct behavioral measurements of impulsivity. Furthermore, despite the adequate sample size (n=300), the number of seropositive participants was relatively low. Therefore, the associations between impulsivity, risky behaviors and infection rates could only be studied using a combined group of seropositive participants. Future studies should explore whether different mechanisms might be involved in different infections and explore generalizability of the current findings to other key populations.

## Conclusions

The present study is an exclusive study examining the role of different aspects of impulsivity in different types of transmission risk behavior among female prisoners in Indonesia. The results show that motor impulsivity (BIS-motor) predicts alcohol-related risk behavior, sensitivity to reward (SPSRQ-reward) predicts drug-related risk behavior, and cognitive impulsivity (BIS-attentional) and goal-directed impulsivity (BAS-drive) predict sexual risk behavior and being seropositive for HIV, HBV, HCV or syphilis. There was no multicollinearity between the predictor variables. The associations between impulsivity and seropositivity were not mediated by risk behavior.

This study shows an association between different types of impulsivity, distinct forms of transmission risk behavior and viral infections (HIV, hepatitis B and C and syphilis). The findings indicate that distinct, independent mechanisms might be involved in different types of risky behavior. Future studies should explore the clinical relevance of these findings for targeted prevention strategies in key populations, such as inmates.

## Acknowledgments

We are grateful to the Jakarta Pondok Bambu Prison for enabling this study. Furthermore, we especially like to thank the prison clinic team: Dr. Jusi, Dr. Nadia and nurse Endah, as well as all other clinical staff who provided us with their help. We also want to thank M. Lenders, N. Kreuzberger, D. Halim and F. Gunawan for collecting the data. Furthermore, we want to thank R. Adiwinata and R. Febrian for preparing the data for further analysis, Prof. L.A. Lesmana and Prof. Z. Djoerban for their advice during the study and A. Rahmalia, M. Herlina Sitinjak and H. Puspa Indah for their help with professional translation of the questionnaires.

